# Adipose tissue macrophages orchestrate β cell adaptation in obesity through secreting miRNA-containing extracellular vesicles

**DOI:** 10.1101/2020.06.12.148809

**Authors:** Hong Gao, Zhenlong Luo, Zhongmou Jin, Yudong Ji, Wei Ying

**Affiliations:** Division of Endocrinology and Metabolism, Department of Medicine, University of California, San Diego, California, USA; Department of Gastroenterology, Tongji Hospital, Tongji medical College, Huazhong University of Science and Technology, Wuhan, China; Division of Biological Sciences, University of California, San Diego, California, USA; Department of Anesthesiology, Tongji Hospital, Tongji medical College, Huazhong University of Science and Technology, Wuhan, China

**Author notes:** To whom correspondence should be addressed: Dr. Wei Ying, Stein Clinical Research Building, Room 229, Department of Medicine, Division of Endocrinology and Metabolism, University of California, San Diego, La Jolla, California, 92093.

**Keywords:** adipose tissue macrophage, extracellular vesicle, miRNA, β cell replication, insulin secretion, obesity

## Abstract

Obesity induces an adaptive expansion of β cell mass and insulin secretion abnormality. Here, we explore a novel role of adipose tissue macrophages (ATMs) in mediating obesity-induced β cell function and proliferation through releasing miRNA-containing extracellular vesicles (EVs). ATM EVs derived from obese mice notably suppress insulin secretion in both *in vivo* and *in vitro* experiments, whereas there are more proliferating β cells in the islets treated with obese ATM EVs. Depletion of miRNAs blunts the ability of obese ATM EVs to regulate β cell responses. miR-155, a highly enriched miRNA within obese ATM EVs, exerts profound regulation on β cell functions, as evidenced by impaired insulin secretion and increased β cell proliferation after miR-155 overexpression in β cells. By contrast, knockout of miR-155 can attenuate the regulation of obese ATM EVs on β cell responses. We further demonstrate that the miR-155-*Mafb* axis plays a critical role in controlling β cell responses. Taken together, these studies show a novel mechanism by which ATM-derived EVs act as endocrine cargoes delivering miRNAs and subsequently mediating β cell adaptation and functional dysfunction in obesity.

---

The prevalence of Type 2 diabetes mellitus (T2DM) has risen dramatically in the past couple of decades(1). Insulin resistance is a central etiologic defect of T2DM(2, 3). Chronic low-grade tissue inflammation, accompanied by an increase in immune cell infiltration, is a hallmark of obesity in both humans and rodents and an important contributor to the pathogenesis of insulin resistance and metabolic diseases(4, 5). Insulin resistance results in a compensatory growth of pancreatic β cells, leading to a status of hyperinsulinemia. Many of obese individuals are prediabetic, in that they will eventually develop T2DM characterized by an insufficient insulin production for insulin resistance.

Adipose tissue is one of the most expanded metabolic organs in obesity, concomitant with a state of chronic and unresolved inflammation(4, 6-10). Numerous studies in both humans and rodents have demonstrated that a remarkable accumulation of proinflammatory macrophages is one of the striking components of obesity-induced adipose tissue inflammatory responses(11-16). These proinflammatory macrophages residing in obese adipose tissues (ATMs) are one of the main drivers for the pathogenesis of obesity-induced tissue inflammation and insulin resistance(17-21). Earlier studies suggested that obese ATM-derived proinflammatory cytokines, such as tumor necrosis factor alpha (TNFα), directly dampen insulin sensitivity(12, 20, 22). However, there was limited improvement on insulin resistance and glucose metabolism in obese human with anti-TNFα antibody therapies(23-26). In addition to the production of proinflammatory cytokines, recently we have shown that obese ATMs can reduce peripheral insulin sensitivity by releasing extracellular vesicle/microRNAs (EV/miRNAs) locally or into the circulation(27). Among the exosomal miRNAs derived from obese ATMs, we have demonstrated the pathogenic effects of miR-155 on cellular insulin action.

Obesity-induced insulin resistance provokes an adaptive expansion of β cell mass, which compensate by secreting increased amount of insulin(28-30). The mechanisms underlying obesity-induced β cell replication have been implicated, including increased plasma glucose concentrations, insulin, expression of hepatocyte growth factors, and islet macrophage-mediated islet inflammation(31-37). Furthermore, previous studies have shown the critical roles of key transcription factors or microRNAs (miRNA) in regulating the genetic network associated with β cell insulin secretion and proliferation in the context of obesity(38). For example, a previous study reported the critical role of miR-155 in facilitating the growth of β cell mass in the context of hyperlipidemia by directly repressing expression of v-maf musculoaponeurotic fibrosarcoma oncogene family, protein B (*Mafb*)(39). Our previous study has demonstrated that obese ATMs preferentially secreted miR-155 enriched EVs, leading to an accumulation of this miRNA in the target cells resided locally or at distal sites(27). However, whether obese ATM-secreted miRNA-containing EVs are capable of regulation of β cell functions in response to obesity remains unknown.

Here, we report that obese ATM-produced EVs can be delivered into β cells, both *in vitro* and *in viv*o, resulting in blunted insulin secretion and more proliferating β cells. The ability of obese ATM EVs to mediate β cell responses are mainly dependent on miRNAs, as evidenced by modest effects of obese DicerKO ATM EVs on β cell insulin secretion and replication. We further identify that miR-155, a highly enriched miRNA within obese ATM EVs, can impair the glucose-stimulated insulin secretion of β cells by suppressing *Mafb* expression. In addition, the miR-155-*Mafb* axis can promote β cell proliferation. By contrast, depletion of miR-155 can partially prevent the effects of obese ATM EVs on β cell responses.

## Research Design and Methods

### Animal

C57BL/6 (B6) mice were fed a high-fat diet (60% fat calories, 20% protein calories, and 20% carbohydrate calories; Research Diets) or a normal chow diet ad libitum. MIPGFP (Stock No. 006864), Mki67 (Stock No. 029802), Dicer flox/flox (Stock No. 006366) and LysMCre mice (Stock No. 004781) were purchased from the Jackson Laboratory. To generate myeloid cell-specific Dicer null mice, Dicer flox/flox were bred with transgenic mice harboring Cre recombinase driven by myeloid-specific lysozyme M promoter to create the following genotypes: Dicer flox/flox (control) and LysMCre-Dicer (DicerKO). To generate MIPGFP-Mki67 mice (GFP+RFP+ cells are proliferating ki67+ β cells.), MIPGFP (GFP+ cells are β cells.) mice were bred with Mki67 (RFP+ cells are proliferating ki67+ cells.) mice. All mice were maintained on a 12/12-hour light-dark cycle.

### Flow cytometry analysis

Unless otherwise specified, we purchased antibodies from Biolegend. Visceral stromal cells were stained with fluorescence-tagged antibodies to macrophages (CD45+CD11b+F4/80+). Data were analyzed using Flowjo software.

### Isolation of adipose tissue macrophages (ATMs)

Visceral adipose tissues (VATs) were mechanically chopped and then digested with collagenase II (Sigma-Aldrich, Cat. No. C2674) for 15 min at 37°C. After passing cells through a 200 µm cell strainer (VWR, Cat. No. 100490-158) and centrifugation at 1,000 x g for 10 min, the pellet containing the stromal vascular cell (SVC) fraction was then incubated with red blood cell lysis buffer. SVC single cell suspensions were incubated with fluorescence-tagged antibodies against CD45 (Biolegend, Cat. No. 103116), CD11b (Biolegend, Cat. No. 101206), and F4/80 (Biolegend, Cat. No. 123116). CD45+CD11b+F4/80+ macrophages were purified using SONY MA900 flow cytometer (SONY). In addition, cells were stained with CD11c (Biolegend, Cat. No. 117343) and CD206 (Biolegend, Cat. No. 141715) antibodies to measure the levels of M1 and M2 activation. ATMs were then cultured in IMDM containing 10% exosome-free FBS to produce extracellular vesicles (EVs).

### Isolation of islets

NCD and HFD mice were euthanized, and freshly-prepared collagenase P (Roche, Cat. No. 11249002001) solution (0.5 mg/ml) was injected into the pancreas via the common bile duct. The perfused pancreas was digested at 37°C for 10 min, and the islets were handpicked under a stereoscopic microscope. To sort out GFP+ or GFP+RFP+ β cells, islets were dispersed into single cell suspensions, and then cells with fluorescent reporters were purified using a SONY MA900 flow cytometer (SONY).

### EV purification and characterization

The EVs from ATM culture medium were prepared as previously described(27). After 24 hours culture, debris and dead cells in the ATM medium were removed by centrifugation at 1,000 x g for 10 min and then filtrated through a 0.2 µm filter. The medium was then subjected to ultracentrifugation at 100,000 x g for 4 hours at 4^°^C. After wash with PBS (100,000 x g for 20 min), the EV-containing pellet was resuspended in 1 ml PBS and passed through a 0.2 µm filter to remove large particles. The particle size and concentration of ATM EVs were measured by NanoSight analysis (Malvern Instruments). To monitor EV trafficking, EVs were labeled with PKH26 fluorescent dye using the PKH26 fluorescent cell linker kit (Sigma-Aldrich, Cat. No. PKH26GL-1KT). After PKH26 staining, the EVs were washed with PBS and collected by ultracentrifugation (100,000 x g for 2 hours) at 4°C. Finally, PKH26 labeled EVs were resuspended in PBS.

To analyze the characteristics of macrophage-derived EVs by flow cytometer (MA900, SONY), EVs (5×10^9^) were incubated with latex beads (1×10^8^; ThermoFisher, Cat. No. A37304) at 4^°^C. After overnight incubation, the beads were washed with PBS twice and finally resuspened in 200 µl of PBS.

### In vivo and in vitro EV treatment

For *in vitro* GSIS assays, 0.2×10^8^ EVs on the basis of NanoSight analysis were added to 10 islets for 24 hours. For *in vitro* proliferation assays, 50 MIPGFP-Mki67 islets were treated with 1×10^8^ EVs for 72 hours. For *in vivo* treatment, 7 weeks old recipient mice were intravenously injected 1×10^9^ EVs twice per week.

### Differentiation of bone marrow-derived macrophages

Bone marrow-derived macrophages (BMDMs) were prepared as previously described(40).

### Depletion of tissue macrophages

To deplete tissue macrophages, WT mice were given clodronate liposomes (80 mg/kg body weight; FormuMax, Cat. No. F70101C-N) every 7 days via intraperitoneal injection(15). These mice also started feeding HFD. The HFD-fed control mice were treated with empty liposomes.

### Transfection of miR-155 mimics, miR-155 hairpin inhibitor, or siRNA

Cy3 labeled miR-155 mimics (GE Dharmacon), miR-155 mimics (ThermoFisher Scientific, Cat. No. 4464066), miR-155 hairpin inhibitor (Horizon, Cat. No. IH-310430-08-0002), or siRNA-*Mafb* (Horizon, Cat. No. J-059035-09-0002) were transfected into recipient cells with the lipofectamine RNAiMAX reagent (ThermoFisher Scientific, Cat. No. 13778-075). After 24 hours, the transfection efficiencies were validated by qPCR analysis.

### Co-culture assay

After transfected with Cy3-miR-155 mimics (GE Dharmacon), obese ATMs (0.1×10^5^/well) were co-cultured with Min6 cells at a ratio of 1:5 using a Transwell plate (0.4 µm polycarbonate filter, Corning) for 16 hours, with Min6 cells placed in the lower chamber and obese ATMs in the upper chamber. After washing with PBS twice, the intensity of Cy3 red fluorescence in Min6 cells was examined by flow cytometer (MA900, SONY). To inhibit EV secretion, obese ATMs containing Cy3-miR-155 were pretreated with GW4869 (an inhibitor of neutral sphingomyelinase, 10 µM; Cayman Chemical, Cat. No. 6823-69-4) for 24 hours(41). These obese ATMs were used to co-cultured with Min6 cells in the medium containing GW4869 for anther 16 hours.

### Immuno-fluorescence staining

Pancreases were cut and snap frozen in optimum cutting temperature (O.C.T., Fisher Healthcare). Six µm cryo-sections of tissue sections were cut and fixed with pre-cold acetone for 20 min. Immunostaining was performed as previously described(36). Before adding insulin antibody (ThermoFisher Scientific, Cat. No. 53-9769-80), sections were blocked with 5% normal donkey serum (Jackson ImmunoResearch, Cat. No. 017-000-001). Nuclei were stained with DAPI (4’,6-Diamidino-2-28 phenylindole dihydrochloride). After washing, slides were incubated with Antifade Mounting Medium (VECTASHIELD, Cat. No. ZF0612) and covered with coverslip. Images were acquired on Keyence Fluorescent Microscope and were processed with ImageJ (NIH).

### Quantitative Reverse Transcriptase-polymerase Chain Reaction (RT-PCR) Analysis

Total RNA was extracted from either islets or GFP+ β using the RNA extraction protocol according to the manufacturer’s instructions (Zymo Research, Cat. No. R1051). cDNA was synthesized using SuperScript III and random hexamers. qPCR was carried out in 10 μl reactions using iTaq SYBR Green supermix on a StepOnePlus Real-Time PCR Systems (ThermoFisher Scientific). The data presented correspond to the mean of 2^-ΔΔCt^ from at least three independent experiments after being normalized to β-actin.

### Glucose-stimulated insulin secretion (GSIS) assays

Primary mouse islets isolated from both lean and obese mice were also used to evaluate the effects of macrophage EVs on GSIS. Static GSIS experiments were done as described previously(36). Briefly, after washing twice with 1g/L glucose DMEM, islets were incubated overnight in 1g/L glucose DMEM, at 37°C, 5% CO_2_. Next day, the islets were washed with fresh 2.8 mM glucose Krebs Ringer Bicarbonate buffer (KRB buffer; 2.6 mM CaCl_2_/2H_2_O, 1.2 mM MgSO_4_/7H_2_O, 1.2 mM KH_2_PO_4_, 4.9 mM KCl, 98.5 mM NaCl, and 25.9 mM NaHCO_3_, supplemented with 20 mM HEPES and 0.2% BSA), and then fasted with 2.8 mM glucose KRB buffer for 30 min. Next, islets were incubated for 60 min in 2.8 mM or 16.3 mM glucose KRB buffer. Insulin concentrations in the supernatant were determined using Ultrasensitive mouse insulin ELISA kits (Alpco, Cat. No. 80-INSHU-E01.1). To calculate the relative insulin secretion for each group, the insulin levels of 2.8 mM glucose treatment were used as the basal insulin, and 16.3 mM glucose-stimulated insulin levels were normalized to the basal insulin.

### Statistics

Tests used for statistical analyses are described in the figure legends. To assess whether the means of two groups are statistically different from each other, unpaired two-tailed Student’s t test was used for statistical analyses using Prism8 software (GraphPad software v8.0; Prism, La Jolla, CA). *P* values of 0.05 or less were considered to be statistically significant. Degrees of significance were indicated in the figure legends. For the results of glucose and insulin tolerance tests, statistical comparisons between two groups at each time point were performed with unpaired two-tailed Student’s t test.

### Study approval

All animal procedures were done in accordance with University of California, San Diego Research Guidelines for the Care and Use of Laboratory Animals and all animals were randomly assigned to cohorts when used.

## Results

### Adipose tissue macrophage-produced EVs are pathogenic for insulin secretion in obesity

We first examined if ATMs (F4/80+CD11b+) can release EVs harboring miRNAs that are then delivered into β cells. After transfection of red fluorescent Cy3-labeled miR-155 mimics, obese ATMs secreted EVs containing Cy3-miR-155 mimics, as shown by the presence of a robust Cy3 fluorescence in these obese ATM EVs (Fig. 1A). In addition, flow cytometry analysis showed that a robust red fluorescent Cy3 signal was detected in GFP+ β cells after the MIPGFP islets cocultured with Cy3-miR-155 containing ATM EVs, suggesting that ATM-derived miRNA-containing EVs can be taken up into β cells (Fig. 1B). Concomitantly, miR-155 was highly enriched in these recipient cells cocultured with obese ATM EVs (Fig. 1C). Furthermore, prior treatment of obese ATMs with an EV secretion inhibitor GW4869 remarkably reduced EV production and the delivery of Cy3-miR-155 from ATMs into Min6 cells in a Transwell (pore size=0.4 µm) coculture, demonstrating that EVs are important cargos extracellularly transporting miRNAs (Fig. 1D). Overall, our results show that ATM-derived miRNA-containing EVs can be taken up into pancreatic β cells.

**Figure 1.**
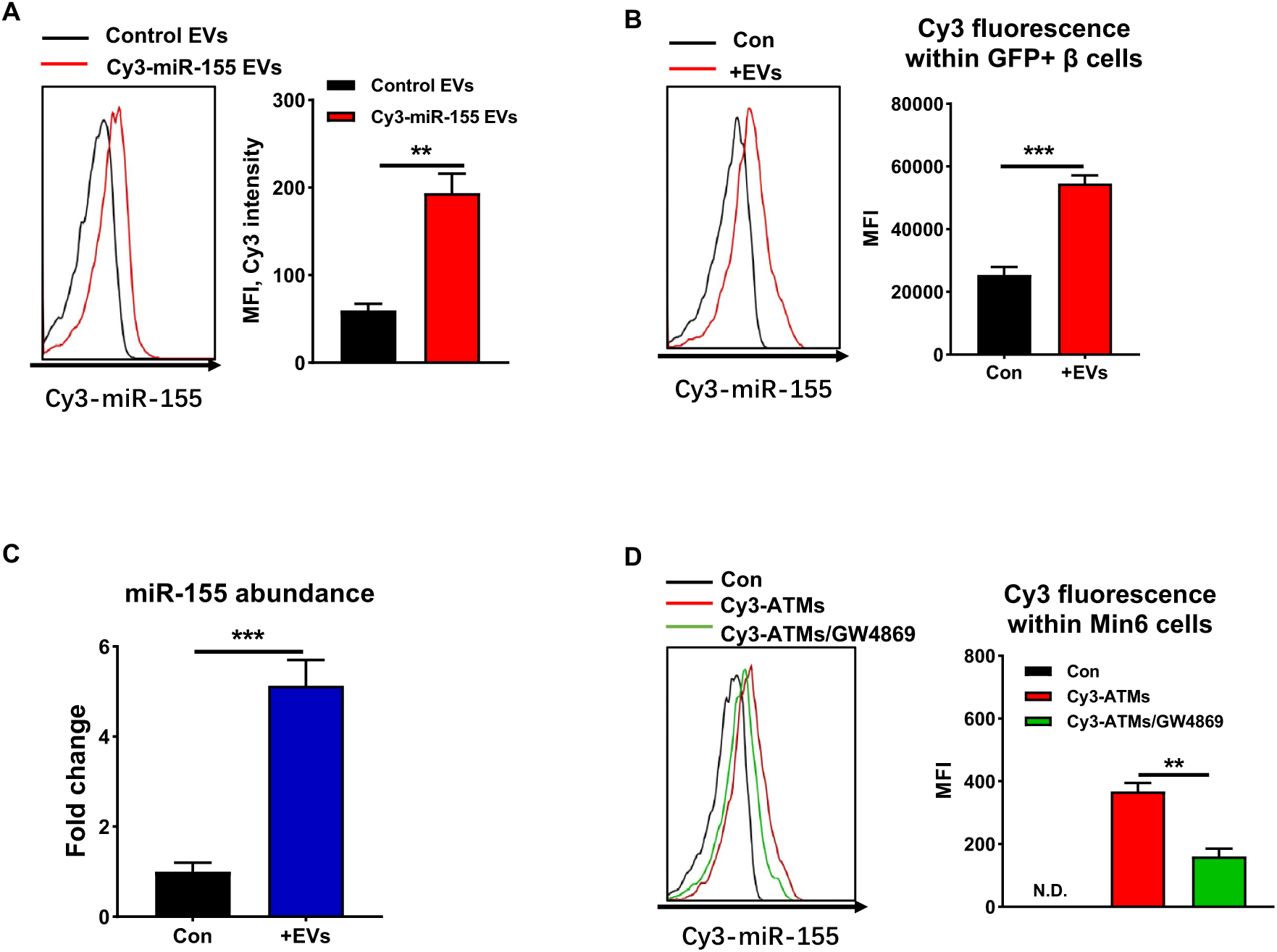
Obese adipose tissue macrophages (ATMs) secrete extracellular vesicles (EVs) as cargos delivering miRNAs into β cells. (A) The delivery of Cy3-miR-155 mimics into EVs derived from obese ATMs transfected with Cy3-miR-155 mimics. (B) Cy3 red fluorescent intensity within GFP+ β cells after MIPGFP islets co-cultured with Cy3-miR-155 containing obese ATM EVs. (C) miR-155 abundance in islets co-cultured with obese ATM EVs. (D) Cy3 red fluorescent intensity within Min6 cells co-culutred with Cy3-miR-155 containing obese ATMs in a Transwell plate with EV secretion inhibitor GW4869 (10 µM). Data are presented as the mean ± SEM. A-D, n=4 per group. ** *P*<0.01, *** *P*<0.001, Student’s t test.

Adipose tissue macrophages play critical roles in the pathogenesis of tissue inflammation and insulin resistance in obesity(4). Our previous study has demonstrated the critical role of ATM-derived EV/miRNAs in mediating peripheral insulin sensitivity(27). Thus, to examine the *in vivo* pathogenic effects of obese ATM EV/miRNAs on β cell functions in obesity, obese ATM EVs (1×10^9^ EVs/mouse) were intravenously injected into a group of obese WT mice. Concordant with an efficient delivery into Min6 cells or GFP+ β cells *in vitro*, obese ATM EVs can be readily transported into pancreatic β cells, as evidenced by the appearance of a robust red fluorescence in the pancreatic insulin-producing cells of recipients after an injection of PKH26-labeled obese ATM EVs (Supplementary Fig. 1A). To determine the roles of ATM EVs in regulating β cell responses in obesity, WT recipients were intravenously injected with obese ATM EVs (1×10^9^ EVs/mouse, twice injection per week) and also started feeding high fat diet (HFD). After 4 weeks, all HFD-fed mice had higher population of ATMs and proinflammatory M1-like (F4/80+CD11b+CD206-CD11c+) macrophages than in lean WT mice (Supplementary Fig. 1B). There was no difference in ATM population and activation among all HFD-fed mice (Supplementary Fig. 1B). In line with our previous findings, the recipients injected with obese ATM EVs exhibited worse obesity-induced glucose intolerance and insulin resistance, as measured by glucose and insulin tolerance tests (Fig. 2A and 2B). Compared to the controls, obese ATM EV treatment resulted in significantly lower insulin levels at feeding condition (Fig. 2C). In addition, obese ATM EVs-treated mice failed to produce additional insulin after glucose injection, whereas control obese mice elevated insulin secretion in response to glucose stimulation (Fig. 2D). Consistently, islets isolated from obese ATM EVs-treated recipients had notably less abundance of genes associated with insulin secretion than did the control islets (Fig. 2E). Concomitant with obese ATM EVs-induced reduction in circulating insulin, these obese mice had less body weight due to a blunted adipose tissue expansion after 4 weeks HFD feeding (Fig. 2F). qPCR analysis also indicated that obese ATM EV treatment resulted in a significant repression on the expression of genes associated with lipogenesis (Supplementary Fig. 1C). We further confirmed the suppressive effect of obese ATM EVs on β cell insulin secretion, as shown by the blunted glucose-stimulated insulin secretion (GSIS) of WT islets after *in vitro* treatment of obese ATM EVs (0.2×10^8^ EVs/10 islets) (Fig. 2G and 2H). Overall, these results suggest that obese ATMs can impair the insulin secretion of β cells through releasing pathogenic EVs.

**Figure 2.**
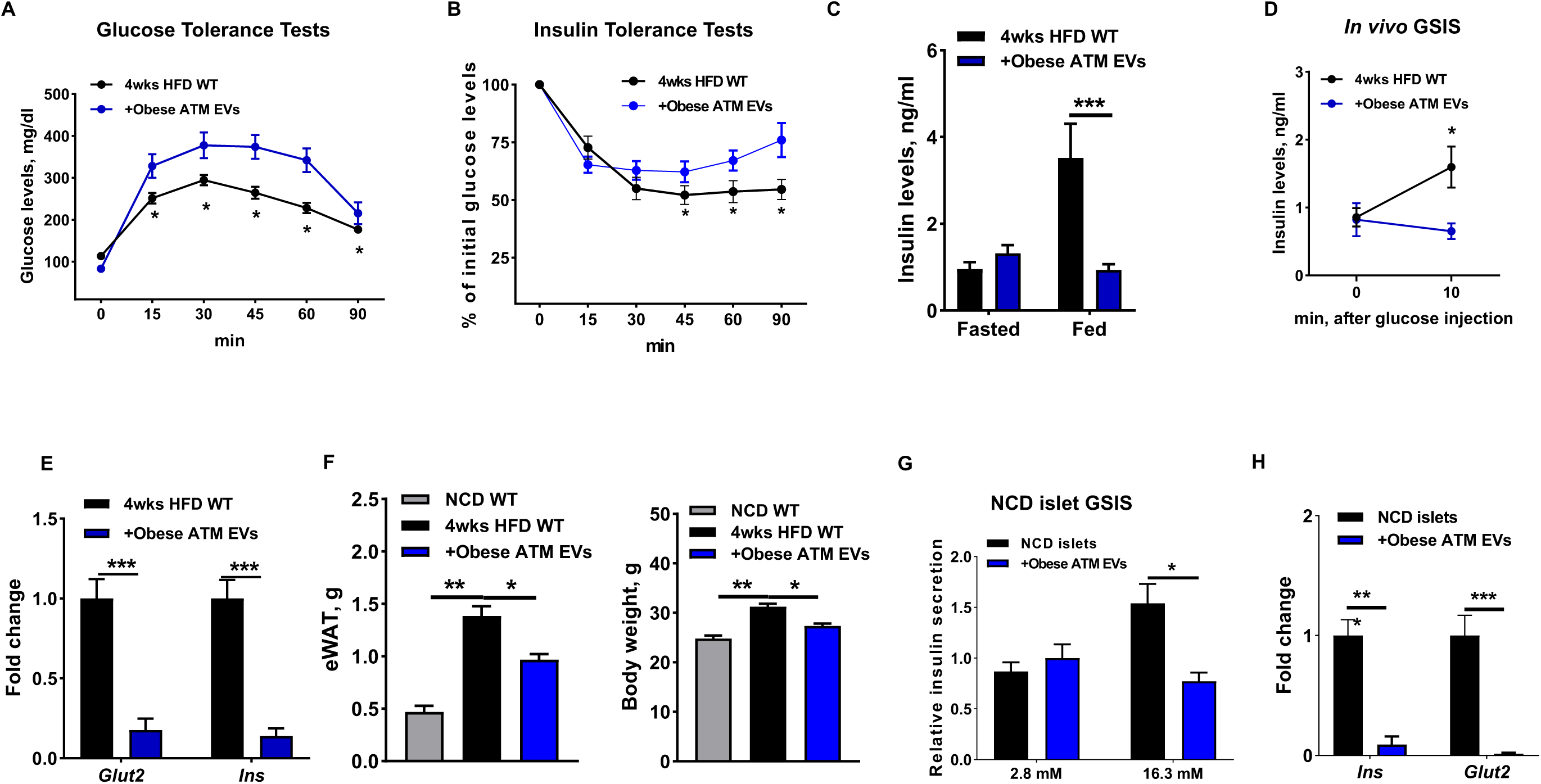
Obese ATM EVs blunt the insulin secretion of obese mice. (A and B) Glucose and insulin tolerance tests were performed in high fat diet (HFD)-fed WT recipient mice after 4 weeks obese ATM EV treatment. (C) Insulin levels after 16 hours fasting or feeding. (D) The insulin levels of HFD recipient mice after 10 min glucose injection. (E) The abundance of genes associated with insulin secretion in the islets of HFD recipients. (F) The epididymal fat mass and body weight of HFD mice after 4 weeks obese ATM EV treatment. (G) The effect of obese ATM EVs on the glucose-stimulated insulin secretion (GSIS) of islets isolated from normal chow diet (NCD) fed mice. (H) The expression of *Ins* and *Glut2* in NCD islets treated with obese ATM EVs. Data are presented as the mean ± SEM. A-F, n=5-6 per group, G and H, n=4 per group. * *P*<0.05, ** *P*<0.01, *** *P*<0.001, Student’s t test.

### Obese ATM-derived EVs can promote obesity-induced β cell proliferation

Obesity-induced insulin resistance induces an expansion of pancreatic β cell mass. To examine the mechanism by which obese ATM-derived EVs enhance β cell proliferation in the context of obesity, obese ATM EVs (1×10^9^ EVs/mouse, twice injection per week) were intravenously injected into a group of HFD-fed MIPGFP-Mki67 mice which harbor the markers labelling proliferating ki67+ β cells (GFP+RFP+). After 4 weeks HFD feeding, the control MIPGFP-Mki67 mice had a modest increase in the proportion of proliferating ki67+ β cells compared to the lean MIPGFP-Mki67 mice (Fig. 3A). Notably, HFD-fed MIPGFP-Mki67 mice injected with obese ATM EVs displayed a significant enhancement in proliferating ki67+ β cell proportion, compared to the control obese MIPGFP-Mki67 mice (Fig. 3A). We also observed an increase in β cell proliferation after 72 hours treatment of NCD MIPGFP-Mki67 islets with obese ATM EVs (1×10^8^ EVs/50 islets) (Fig. 3B). qPCR analysis also confirmed that the expression of genes associated with β cell mass expansion were remarkably increased after obese ATM EV treatment (Fig. 3C). Thus, these data indicate an improvement of obese ATM EVs on β cell proliferation.

**Figure 3.**
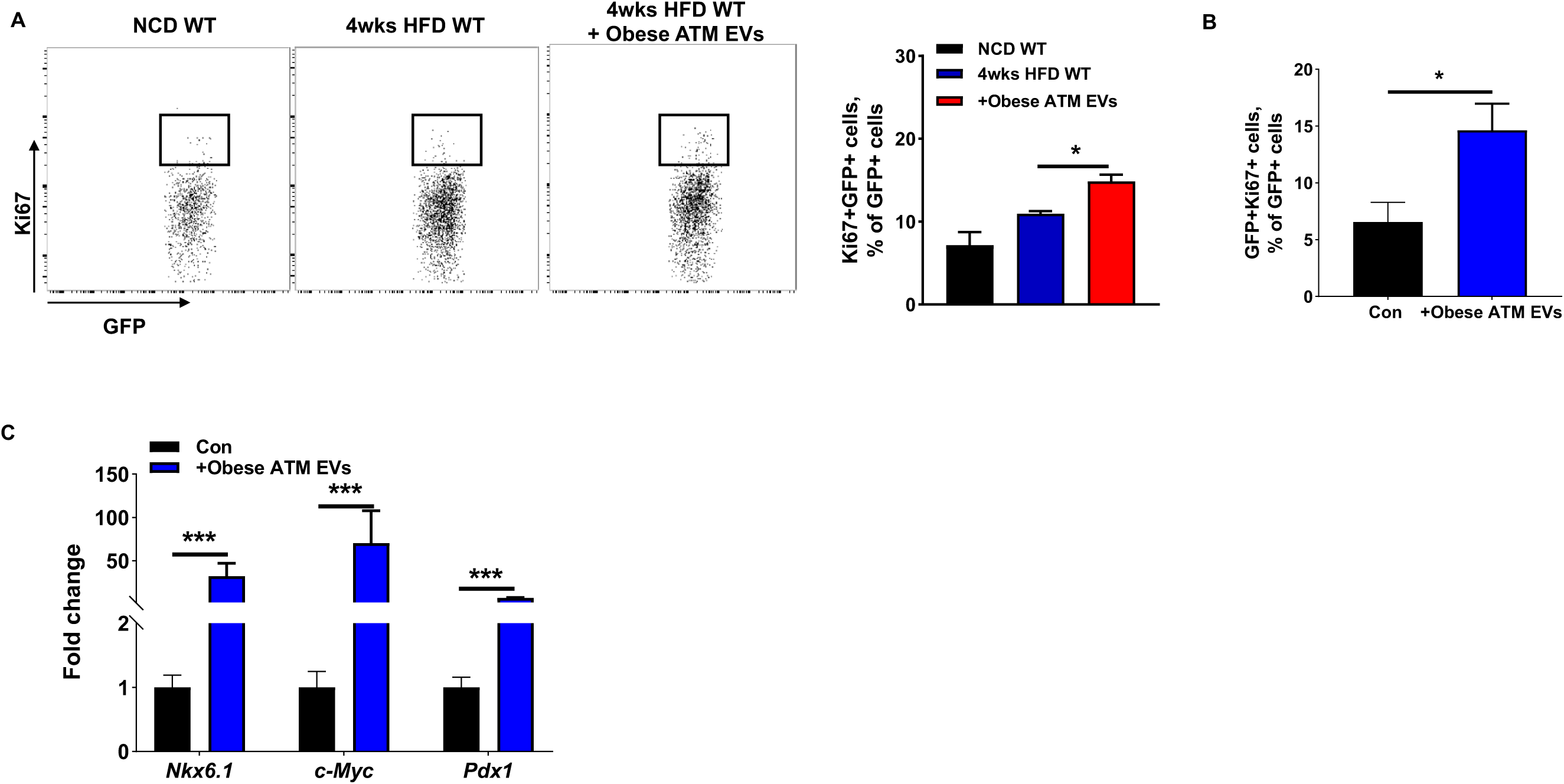
Obese ATM EVs enhance β cell proliferation in response to obesity. (A) The population of Ki67+GFP+ cells in the islets isolated from NCD MIPGFP-Mki67 WT, 4wks HFD MIPGFP-Mki67 WT, or 4wks HFD MIPGFP-Mki67 WT mice treated with obese ATM EVs. (B) The *in vitro* effect of obese ATM EVs on the Ki67+GFP+ cell proportion of NCD MIPGFP-Mki67 islets. (C) The abundance of genes associated with β cells proliferation after 72 hours obese ATM EV treatment. Data are presented as the mean ± SEM. A, n=5 per group; B and C, n=4 per group. * *P*<0.05, *** *P*<0.001, Student’s t test.

### miRNAs are indispensable components for the regulation of obese ATM-derived EVs on β cell responses

To demonstrate that obese ATM EVs-induced β cell responses are dependent on miRNA functions, we collected miRNA-free EVs from obese ATMs without Dicer which is an essential ribonuclease for the production of mature miRNAs (Supplementary Fig. 2A)(42). qPCR results confirmed the absence of miRNAs in obese DicerKO ATMs, as measured by the abundance of miR-155 that is highly expressed in obese ATMs (Supplementary Fig. 2B). Consistently, treatment of β cells with obese ATM EVs led to impaired GSIS, whereas WT islets treated with miRNA-free EVs had comparable levels of insulin secretion after glucose stimulation with the control cells (Fig. 4A). Additionally, mice treated with miRNA-free EVs had comparable metabolic phenotypes with the controls, as measured by fat mass, body weight, GTT, ITT, insulin levels, and GSIS (Fig. 4B-4G). HFD-fed MIPGFP-Mki67 mice treated with miRNA-free EVs had a comparable population of GFP+RFP+ proliferating β cells with the control obese mice (Fig. 4H). These results indicate that miRNAs are important components which contribute to the ability of obese ATM EVs to regulate peripheral insulin sensitivity and β cell functions.

**Figure 4.**
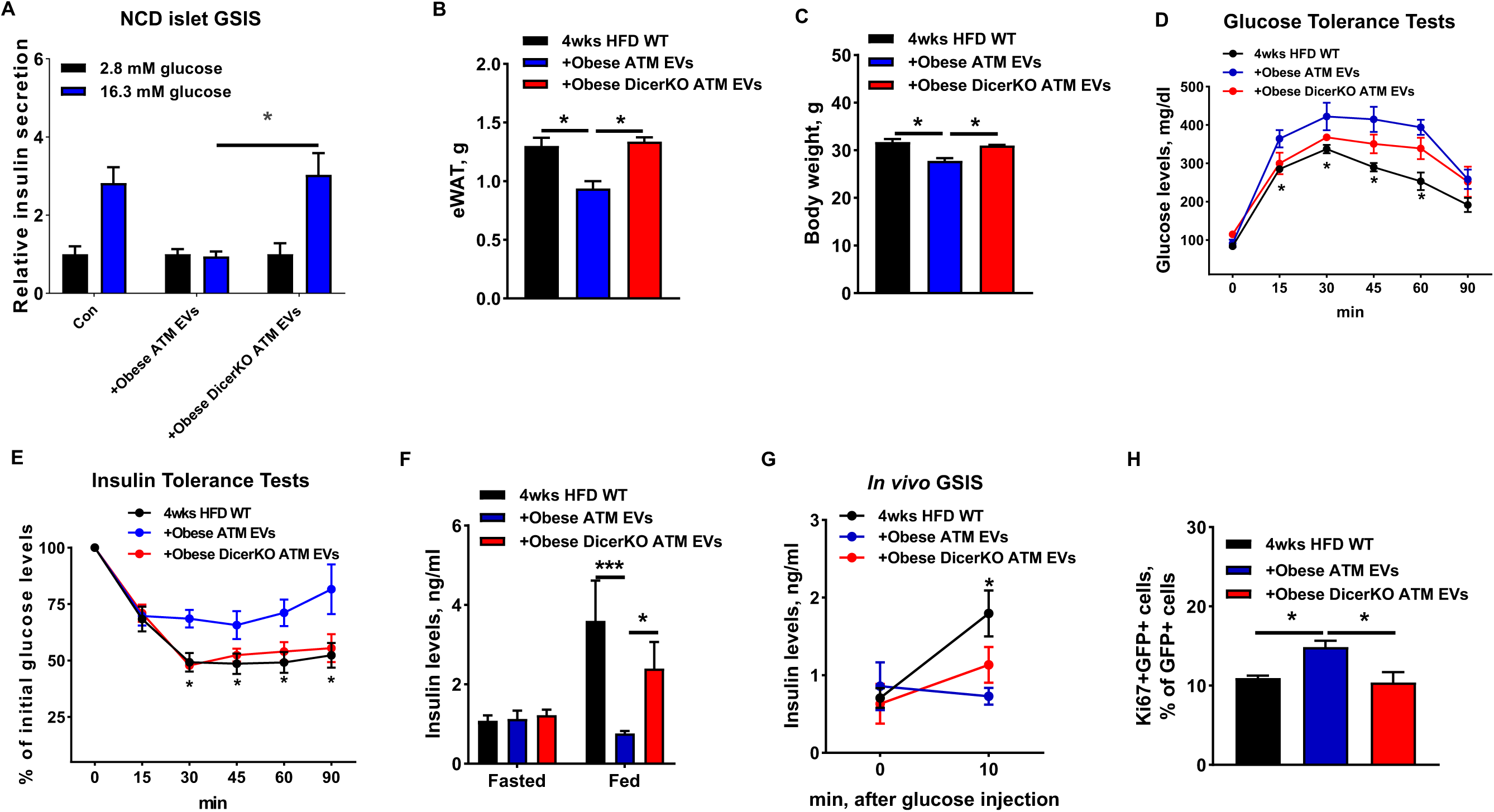
The importance of miRNAs in obese ATM EVs. (A) The GSIS of NCD islets after either obese ATM EV or obese DicerKO ATM EV treatment. After 4 weeks obese ATM EV or obese DicerKO ATM EV treatment, the epididymal fat mass (B), body weight (C), glucose and insulin tolerance tests (D and E), insulin levels after 16 hours fasting or feeding (F), and glucose-stimulated insulin levels (G) of HFD recipients were measured. (H) The effect of obese ATM EVs on Ki67+GFP+ cell population of 4wks HFD MIPGFP-Mki67 mice. Data are presented as the mean ± SEM. A, n=4 per group; B-H, n=5 per group. * *P*<0.05, Student’s t test.

### miR-155 is a functional molecule within obese ATM EVs mediating β cell responses

Given the critical roles of miRNAs in obese ATM EVs, we next explored the mechanisms by which obese ATM-derived EV miRNAs regulate β cell responses. Majority of obese ATMs are proinflammatory macrophages that preferentially produce miR-155, and our previous study has revealed that miR-155 is one of the highly enriched miRNAs in obese ATM EVs(27, 43). We also confirmed that the GFP+ β cells of recipients accumulated more miR-155 after 4 weeks treatment of obese ATM EVs, compared to the controls (Fig. 5A). We next examined the impact of miR-155 on insulin secretion by overexpression of miR-155 in β cells. The lean islets transfected with miR-155 mimics showed a remarkable decrease in insulin secretion in the presence of high glucose concentration (16.3 mM) (Fig. 5B, Supplementary Fig. 3A). The expression of genes associated with insulin secretion were also significantly reduced in the GFP+ β cells with miR-155 overexpression (Fig. 5C). Interestingly, obesity resulted in more miR-155 accumulated in β cells, as evidenced by a higher miR-155 abundance in the GFP+ β cells isolated from 8wks HFD MIPGFP mice than in the β cells of lean MIPGFP mice (Fig. 5D). 4wks HFD feeding led to a modest increase in β cells miR-155 abundance (Supplementary Fig. 3B). Notably, repression of miR-155 activity by a miRNA hairpin inhibitor transfection can restore the GSIS of obese islets (Fig. 5E). In contrast to the suppressive effect on insulin secretion, miR-155 overexpression resulted in a higher percentage of proliferating ki67+ β cells (GFP+RFP+) in islets isolated from NCD MIPGFP-Mki67 mice (Fig. 5F). Thus, these data suggest that amassing miR-155 in β cells can impair insulin secretion but promote β cell proliferation.

**Figure 5.**
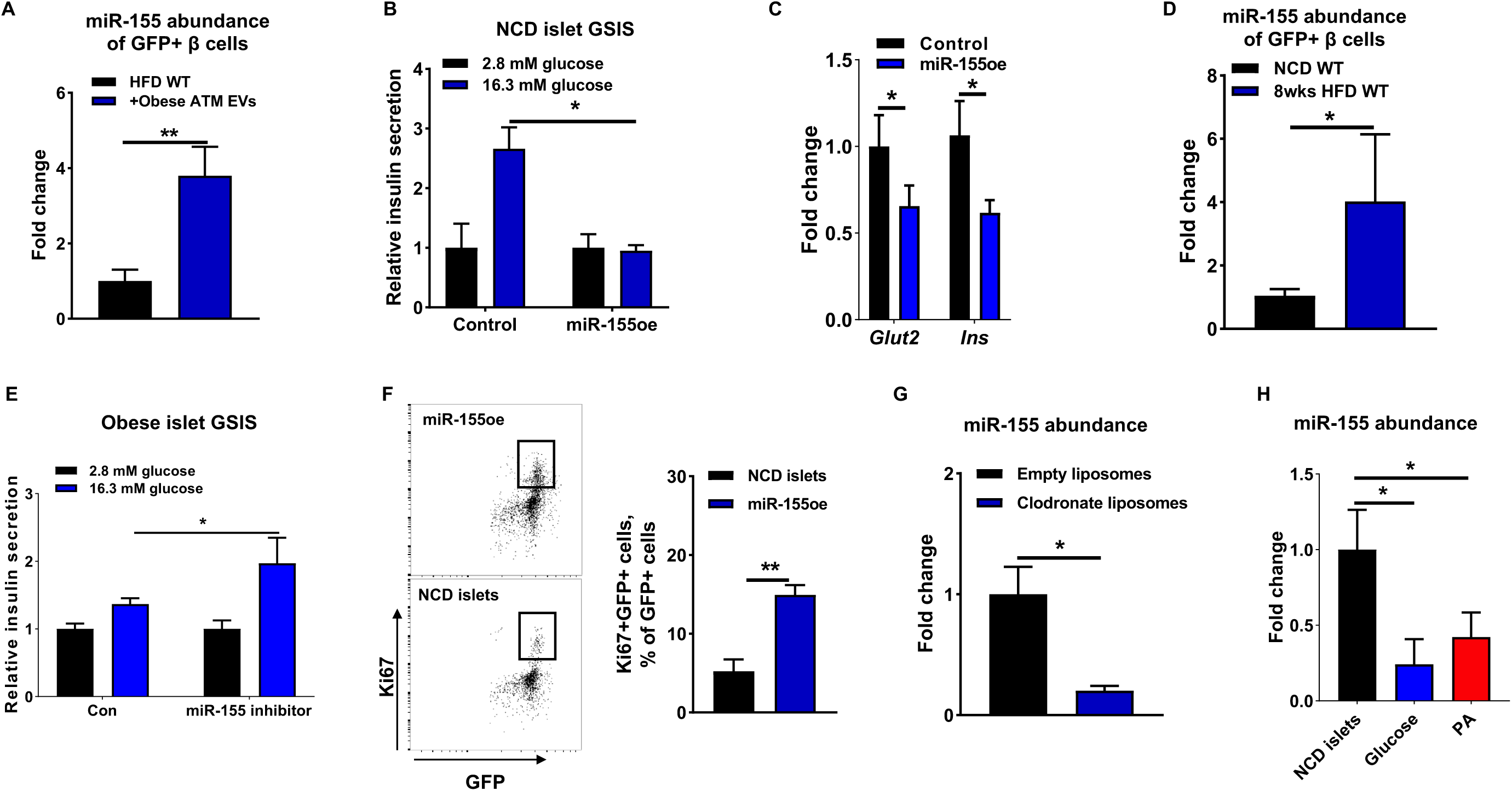
miR-155 accumulation leads to a reduction in insulin secretion but an increase in β cell proliferation. (A) miR-155 abundance of GFP+ cells after HFD MIPGFP-Mki67 mice treated with obese ATM EVs for 4 weeks. (B and C) The effect of miR-155 overexpression (miR-155oe) on the GSIS and genes associated with insulin secretion of NCD islets. (D) miR-155 abundance of GFP+ cells isolated from NCD or 8 wks HFD MIPGFP-Mki67 mice. (E) The GSIS of obese islets after transfection of miR-155 inhibitor. (F) The proportion of Ki67+GFP+ cells in NCD MIPGFP-Mki67 islets transfected with miR-155 mimics. (G) miR-155 abundance in islets of obese mice treated with either empty liposomes or clodronate liposomes. (H) The effect of glucose (16.3 mM) or palmitate acid (PA; 500 µM) on miR-155 levels in NCD islets. Data are presented as the mean ± SEM. A-H, n=4 per group. * *P*<0.05, ** *P*<0.01, Student’s t test.

Obesity switches macrophage activation from anti-inflammation to proinflammatory status, preferentially secreting miR-155(27). Thus, we next determined the contribution of macrophages to miR-155 abundance in target cells by the clodronate liposome-mediated depletion of macrophages (Supplementary Fig. 3C). Notably, depletion of macrophages in obese WT mice resulted in less miR-155 abundance in islets, compared to the obese control treated with empty liposomes (Fig. 5G). In addition, to examine the regulation of hyperglycemia or hyperlipidemia on islet miR-155 abundance, islets isolated from lean WT mice were treated with glucose (16.3 mM) or palmitate acid (500 µM) *in vitro*. After 24 hours treatment, miR-155 expression was remarkably suppressed in these islets, compared to the control cells (Fig. 5H). Therefore, these data suggest that macrophages are the major contributors to the accumulation of miR-155 in islets in obesity.

To further validate that miR-155 is one of the functional miRNAs contributing the ability of obese ATM EVs to mediate β cell functions, we collected the miR-155 enriched EVs from the DicerKO bone marrow-derived macrophages (BMDMs; lack of mature miRNAs) transfected with miR-155 mimics (Supplementary Fig. 4A). We confirmed that the transfected miR-155 mimics can be packed into BMDM-derived EVs, as evidenced by a strong Cy3 red fluorescence detected in the EVs secreted from the BMDMs overexpressed with Cy3-miR-155 (Supplementary Fig. 4B). In addition, we observed the appearance of Cy3 red fluorescence in pancreas after intravenous injection of Cy3-miR-155 enriched BMDM EVs (Supplementary Fig. 4C). qPCR analysis also indicated an efficient delivery of miR-155 into GFP+ β cells after treatment of HFD-fed MIPGFP-Mki67 mice with miR-155-enriched BMDM EVs for 4 weeks (Fig. 6A). Compared to the control mice, 4wks HFD-fed MIPGFP-Mki67 recipients showed impaired glucose tolerance, insulin sensitivity, insulin levels, and *in vivo* GSIS after 4 weeks injection of miR-155-enriched BMDM EVs (Fig. 6B-6E). These mice treated with miR-155-enriched BMDM EVs had a significant reduction in fat mass and body weight, compared to the control mice (Fig. S4D and S4E). Additionally, there was an increased population of proliferating ki67+ β cells (GFP+RFP+) after 4 weeks miR-155-enriched BMDM EV treatment (Fig. 6F). Consistent with the suppressive effect of miR-155-enriched macrophage EVs on *in vivo* GSIS, miR-155-enriched BMDM EV treatment resulted in decreased glucose-stimulated insulin secretion, compared to the control cells (Fig. 6G).

**Figure 6.**
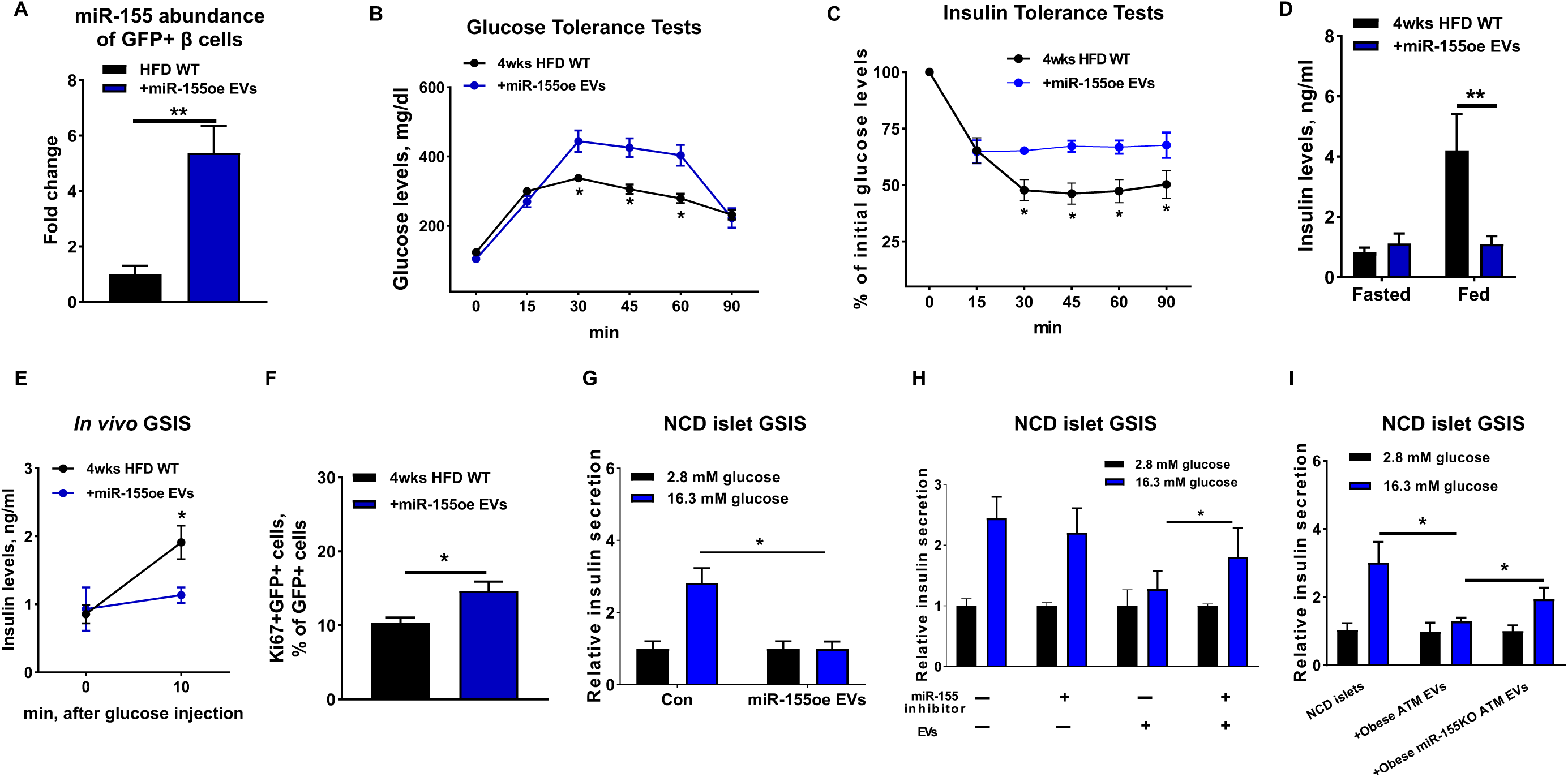
miR-155 is a key molecule contributing to the ability of obese ATM EVs to regulate β cell responses. After 4 weeks treatment of miR-155 enriched BMDM EVs (miR-155oe EVs), the miR-155 expression of GFP+ cells (A), glucose and insulin tolerance tests (B and C), insulin levels after 16 hours fasting or feeding (D), and glucose-induced insulin levels (E) of HFD MIPGFP-Mki67 recipients were measured. (F) The population of ki67+GFP+ cells in the islets of HFD MIPGFP-Mki67 mice after 4 weeks treatment of miR-155oe EVs. (G) The effect of miR-155oe EVs on NCD islet GSIS. (H) The GSIS of NCD islets treated with miR-155 inhibitor or obese ATM EVs. (I) The GSIS of NCD islets treated with EVs derived from either obese ATMs or obese miR-155KO ATMs. Data are presented as the mean ± SEM. A-F, n=5 per group; G-I, n=4 per group. * *P*<0.05, ** *P*<0.01, Student’s t test.

We next evaluated the impact of miR-155 depletion on the ability of obese ATM EVs to blunt insulin secretion. Treatment of miR-155 inhibitor partially restored the GSIS of islets exposed to obese ATM EVs (Fig. 6H). We also collected the miR-155 free EVs from obese miR-155KO ATMs. Depletion of miR-155 from ATM EVs alleviated the pathogenic effect of obese ATM EVs on insulin secretion, as evidenced by a greater GSIS of islets treated with miR-155-free EVs than those treated with obese ATM EVs (Fig. 6I). However, after glucose stimulation, miR-155 free EV-treated islets produced less insulin than the control cells, suggesting miR-155 is likely not the only pathogenic molecule in obese ATM EVs (Fig. 6I). Taken together, these results demonstrate that miR-155 is one of the key functional molecules contributing to the ability of obese ATM EVs to regulate β cell functions.

### The miR-155-*Mafb* axis exerts profound regulation on the insulin secretion and proliferation of β cells

It has been demonstrated that miR-155 regulates β cell functions through suppressing the expression of *Mafb(39)*. Consistent with these previous reports, miR-155 accumulation in β cells was accompanied by a significant decrease in *Mafb* abundance (Fig. 7A, Supplementary Fig. 5A and 5B). We further confirmed the critical role of *Mafb* in facilitating β cell insulin secretion, as shown by a significant reduction in GSIS after a short interfering RNA-induced *Mafb* downregulation (Fig.7B and 7C, Supplementary Fig. 5C). By contrast, compared to the control MIPGFP-Mki67 islets, knockdown of *Mafb* led to an increase in the population of GFP+RFP+ proliferating ki67+ β cells (Fig. 7D). By contrast, suppression of miR-155 function by a miRNA hairpin inhibitor transfection can restore the expression level of *Mafb* in β cells treated with either obese ATM EVs, accompanied by the recovery of GSIS and β cell proliferation (Fig. 6H, 7E, and 7F). Thus, these results suggest that the miR-155 regulates β cell responses through a direct inhibition of *Mafb*.

**Figure 7.**
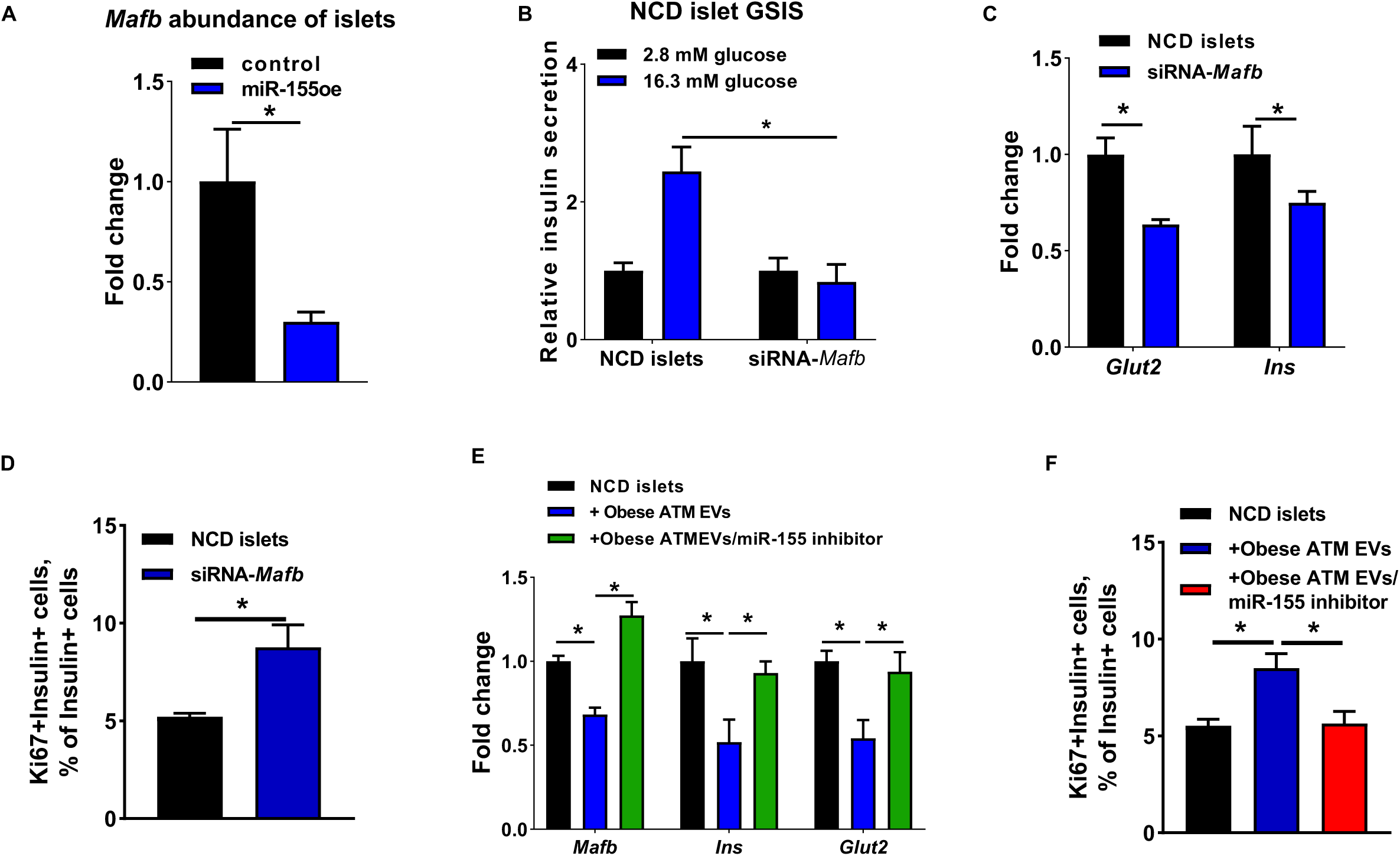
The miR-155-*Mafb* axis exerts profound regulation on β cell functions. (A) The *Mafb* abundance of islets transfected with miR-155 mimics. (B and C) The GSIS and expression of *Glut2* and *Ins* after transfection of siRNA-*Mafb* into NCD islets. (D) The population of Ki67+GFP+ cells in the NCD MIPGFP-Mki67 islets transfected with siRNA-*Mafb*. (E and F) The effects of miR-155 inhibitor on *Mafb, Ins*, and *Glut2* abundance and β cell proliferation of NCD MIPGFP-Mki67 islets treated with obese ATM EVs. Data are presented as the mean ± SEM. A-F, n=4 per group. * *P*<0.05, Student’s t test.

## Discussion

Here, we present a novel mechanism by which obese ATMs exert profound regulation on obesity-induced β cell proliferation and abnormality of insulin secretion through the secretion of miRNA-containing EVs. After delivered into β cells, obese ATM-derived EVs can remarkably blunt insulin secretion in both *in vivo* and *in vitro* experiments, while there was an improvement on the proliferation of β cells treated with obese ATM EVs. miRNAs are the key components within obese ATM EVs, as evidenced by modest effects of obese DicerKO ATM EVs on β cell functions. Among the identified miRNAs within obese ATM EVs, a high abundance of miR-155 contributes to the capacity of obese ATM EVs to regulate β cell functions through directly repressing on *Mafb* expression. Taken together, these findings highlight a novel role of adipose tissue macrophages in modulating β cell adaptive proliferation and insulin secretion dysfunction in obesity.

Emerging evidence, including our previous study, indicates that EVs serve as important vehicles to transfer a variety of functional molecules between adjacent or distant cells(27, 44-46). In the current study, we have shown that obese ATMs released extracellular miRNAs into β cells, as demonstrated by a robust enrichment of Cy3-miR-155 in β cells after either cocultured with the obese ATMs harboring Cy3-miR-155 in a Transwell system or an addition of obese ATM EVs encapsulating Cy3-miR-155. Supplementation of EV secretion inhibitor GW4869 in the Transwell coculture system significantly reduced the amount of Cy3-miR-155 accumulated in Min6 cells, thus demonstrating the critical role of EVs as carriers for obese ATMs-derived extracellular miRNAs. Concordantly, our *in vivo* studies also confirmed the efficient delivery of miRNA-containing obese ATM EVs into pancreatic β cells after an intravenous injection of Cy3-miR-155 containing EVs or PKH26-labeled EVs into WT recipient mice. In current study, we harvested macrophage-derived EVs following the methods described in our previous studies(27), yielding a variety of extracellular vesicles with a diameter of 50-250 nanometers. More importantly, as previously reported, most of these macrophage EVs contain miRNAs, which is concordant with the presence of Cy3 red fluorescence in the EVs secreted by Cy3-miR-155 containing obese ATMs.

Adipose tissue macrophage population dramatically increases in obesity following the expansion of adipose tissues(11). In this study, we have demonstrated the concept that proinflammatory ATMs can remotely control β cell insulin secretion and proliferation in response to obesity by transferring miRNA-encapsulating EVs into β cells. A major piece of evidence for this new role of obese ATM EVs in regulating β cell functions comes from the reconstitution of obese ATMs-secreted EVs *in vivo* by intravenous injection into the recipients starting HFD feeding. In line with previous findings(14, 47), in contrast to the prolonged obesity-induced expansions of proinflammatory M1 macrophages and beta cell populations, 4 weeks HFD feeding led to modest growth rates of β cell mass and proinflammatory M1 ATM population. After elevating the level of obese ATM EVs by multiple intravenous injections, HFD recipient mice had less glucose-stimulated insulin secretion and insulin levels at feeding condition, while the supplementation of obese ATM EVs led to more proliferating ki67+ β cells. Due to the insufficient amount of circulating insulin to stimulate adipogenesis, the obese ATM EVs treated mice strikingly had much less adipose tissue mass and body weight after 4 weeks HFD feeding. Given the magnitude of the effects of obese ATM EVs *in vivo*, we also confirmed the regulation of these EVs on islet functions with the *in vitro* experiments. These phenotypes are consistent with the previous observation that obesity induces the growth of β cell mass but not the capacity of insulin secretion(48), suggesting the important regulation of obese ATM EVs on β cell responses in the context of obesity.

Many of EV functions have been attributed to EV-encapsulated miRNAs. Our results also have demonstrated that miRNAs are important components within obese ATM EVs. miRNA-depleted EVs derived from obese ATMs without Dicer, an essential enzyme for the synthesis of mature miRNAs, had modest effects on insulin secretion and proliferation of β cells in both *in vivo* and *in vitro* experiments. However, these studies can not exclude the potential functions of other EV components such as proteins and lipids which have been reported their profound functions in previous studies(46). In addition, there is no report about the functions of non-miRNA components within ATM EVs.

Our previous study has revealed the profile of miRNA components harbored by obese ATM EVs(27). Among these miRNAs, miR-155 is highly enriched within obese ATM EVs that can efficiently transfer this miRNA into recipient cells. In addition, compared to other cell types, miR-155 is notably expressed in proinflammatory macrophages (http://biogps.org/#goto=genereport&id=100653389)(43). Using a macrophage-depleted mouse model by clodronate liposome treatment, we further validated that macrophages are one of the main sources releasing extracellular miR-155 into β cells, as shown by much less miR-155 abundance in the islets of clodronate liposome-treated obese mice than in the controls. In addition, *in vitro* glucose or palmitate acid treatment repressed the miR-155 expression in islets, in contrast to the observation that miR-155 was highly expressed in obese β cells compared to the β cells isolated from lean WT mice. This suggests that a sizable component of the obesity-induced increase of miR-155 is likely derived from macrophages.

Previous studies have demonstrated the critical roles of miR-155 in various cell types, including immune cells and insulin sensitizing cells(27, 43, 49, 50). In current study, we found that accumulation of miR-155 produces dual effects on β cell insulin secretion and proliferation, as shown by miR-155-mediated reduction in insulin secretion and enhancement on β cell proliferation. Addition of miR-155 inhibitor can prevent the accumulation of miR-155 as a result of obese ATM EVs incorporated into β cells, leading to the recovery of insulin secretion.

miRNAs exert their biological functions by either blocking translation and/or inducing degradation of target mRNAs by base-pairing to recognition sites(42). In line with the profound effects of the miR-155-*Mafb* regulatory axis on β cell proliferation from a previous report(39), we also found that miR-155-mediated *Mafb* repression led to increased number of proliferating ki67+ β cells *in vivo* and *in vitro*. In addition, we found that knockdown of *Mafb* in β cells can blunt glucose-stimulated insulin secretion, accompanied by a significant repression on the expression of *Glut2* that are critical for insulin secretion.

Given the profound regulation of miR-155 on β cell insulin secretion and proliferation, knockout of miR-155 from obese ATM EVs significantly attenuates the effects of these EVs on β cell responses. In addition, reconstitution of miR-155 in the DicerKO macrophage-derived EVs can yield a comparable capacity to regulate β cell function with obese ATM EVs, suggesting miR-155 is an indispensable component in obese ATM EVs. However, in addition to miR-155, it is likely that other miRNAs within obese ATM EVs also exert profound regulation on β cell insulin secretion and proliferation.

In summary, our current studies have established a novel link between ATMs and β cell by secretion of miRNA-containing ATM EVs in the context of obesity. Treatment of obese ATM EVs causes a reduction in both *in vivo* and *in vitro* insulin secretion, while β cell proliferation is elevated by obese ATM EVs. Based on these studies, we suggest that obese ATM EVs play a critical role in the endocrine signaling system which can exert profound regulation on β cell responses in the context of obesity.

## Acknowledgements

We thank Jachelle M. Ofrecio for helping with insulin ELISA assays.

## Funding

This study was funded by the U.S. National Institute of Diabetes and Digestive and Kidney Diseases K99/R00 award (K99DK115998 to W. Y.) and National Natural Science Foundation of China (No. 81500436 to Z. L.).

## Author contributions

WY designed the studies and HG performed most of the experiments. ZL assisted with preparation of cryosections and immunofluorescent staining. ZJ performed bone marrow cell isolation and differentiation. YJ and ZJ assisted with tissue collection. HG and WY analyzed and interpreted the data. WY supervised the project and wrote the manuscript. WY is the guarantor of this work and had full access to all the data in the study and takes responsibility for the integrity of the data and the accuracy of the data analysis.

## Conflict of interest statements

The authors have declared that no conflict of interest exists.

## Notes

### Competing Interest Statement

The authors have declared no competing interest.

## References

1. Cho NH, Shaw JE, Karuranga S, Huang Y, da Rocha Fernandes JD, Ohlrogge AW, et al. IDF Diabetes Atlas: Global estimates of diabetes prevalence for 2017 and projections for 2045. Diabetes Res Clin Pract. 2018;138:271–81.

2. Kahn SE, Hull RL, and Utzschneider KM. Mechanisms linking obesity to insulin resistance and type 2 diabetes. Nature. 2006;444(7121):840–6.

3. Roden M, and Shulman GI. The integrative biology of type 2 diabetes. Nature. 2019;576(7785):51–60.

4. Lee YS, Wollam J, and Olefsky JM. An Integrated View of Immunometabolism. Cell. 2018;172(1-2):22–40.

5. Romeo GR, Lee J, and Shoelson SE. Metabolic syndrome, insulin resistance, and roles of inflammation--mechanisms and therapeutic targets. Arterioscler Thromb Vasc Biol. 2012;32(8):1771–6.

6. Hirosumi J, Tuncman G, Chang L, Gorgun CZ, Uysal KT, Maeda K, et al. A central role for JNK in obesity and insulin resistance. Nature. 2002;420(6913):333–6.

7. Holland WL, Bikman BT, Wang LP, Yuguang G, Sargent KM, Bulchand S, et al. Lipid-induced insulin resistance mediated by the proinflammatory receptor TLR4 requires saturated fatty acid-induced ceramide biosynthesis in mice. J Clin Invest. 2011;121(5):1858–70.

8. Hotamisligil GS, Arner P, Caro JF, Atkinson RL, and Spiegelman BM. Increased adipose tissue expression of tumor necrosis factor-alpha in human obesity and insulin resistance. J Clin Invest. 1995;95(5):2409–15.

9. Shoelson SE, Lee J, and Yuan M. Inflammation and the IKK beta/I kappa B/NF-kappa B axis in obesity- and diet-induced insulin resistance. Int J Obes Relat Metab Disord. 2003;27 Suppl 3:S49-52.

10. Xu H, Barnes GT, Yang Q, Tan G, Yang D, Chou CJ, et al. Chronic inflammation in fat plays a crucial role in the development of obesity-related insulin resistance. J Clin Invest. 2003;112(12):1821–30.

11. Weisberg SP, McCann D, Desai M, Rosenbaum M, Leibel RL, and Ferrante AW, Jr. Obesity is associated with macrophage accumulation in adipose tissue. J Clin Invest. 2003;112(12):1796–808.

12. Lumeng CN, Deyoung SM, Bodzin JL, and Saltiel AR. Increased inflammatory properties of adipose tissue macrophages recruited during diet-induced obesity. Diabetes. 2007;56(1):16–23.

13. Lumeng CN, DelProposto JB, Westcott DJ, and Saltiel AR. Phenotypic switching of adipose tissue macrophages with obesity is generated by spatiotemporal differences in macrophage subtypes. Diabetes. 2008;57(12):3239–46.

14. Lee YS, Li P, Huh JY, Hwang IJ, Lu M, Kim JI, et al. Inflammation is necessary for long-term but not short-term high-fat diet-induced insulin resistance. Diabetes. 2011;60(10):2474–83.

15. Li P, Oh DY, Bandyopadhyay G, Lagakos WS, Talukdar S, Osborn O, et al. LTB4 promotes insulin resistance in obese mice by acting on macrophages, hepatocytes and myocytes. Nat Med. 2015;21(3):239–47.

16. Kanda H, Tateya S, Tamori Y, Kotani K, Hiasa K, Kitazawa R, et al. MCP-1 contributes to macrophage infiltration into adipose tissue, insulin resistance, and hepatic steatosis in obesity. J Clin Invest. 2006;116(6):1494–505.

17. Lumeng CN, Deyoung SM, and Saltiel AR. Macrophages block insulin action in adipocytes by altering expression of signaling and glucose transport proteins. Am J Physiol Endocrinol Metab. 2007;292(1):E166–74.

18. Li P, Liu S, Lu M, Bandyopadhyay G, Oh D, Imamura T, et al. Hematopoietic-Derived Galectin-3 Causes Cellular and Systemic Insulin Resistance. Cell. 2016;167(4):973–84 e12.

19. Han MS, Jung DY, Morel C, Lakhani SA, Kim JK, Flavell RA, et al. JNK expression by macrophages promotes obesity-induced insulin resistance and inflammation. Science. 2013;339(6116):218–22.

20. Uysal KT, Wiesbrock SM, Marino MW, and Hotamisligil GS. Protection from obesity-induced insulin resistance in mice lacking TNF-alpha function. Nature. 1997;389(6651):610–4.

21. Patsouris D, Li PP, Thapar D, Chapman J, Olefsky JM, and Neels JG. Ablation of CD11c-positive cells normalizes insulin sensitivity in obese insulin resistant animals. Cell Metab. 2008;8(4):301–9.

22. Suganami T, Nishida J, and Ogawa Y. A paracrine loop between adipocytes and macrophages aggravates inflammatory changes: role of free fatty acids and tumor necrosis factor alpha. Arterioscler Thromb Vasc Biol. 2005;25(10):2062–8.

23. Dominguez H, Storgaard H, Rask-Madsen C, Steffen Hermann T, Ihlemann N, Baunbjerg Nielsen D, et al. Metabolic and vascular effects of tumor necrosis factor-alpha blockade with etanercept in obese patients with type 2 diabetes. J Vasc Res. 2005;42(6):517–25.

24. Lo J, Bernstein LE, Canavan B, Torriani M, Jackson MB, Ahima RS, et al. Effects of TNF-alpha neutralization on adipocytokines and skeletal muscle adiposity in the metabolic syndrome. Am J Physiol Endocrinol Metab. 2007;293(1):E102–9.

25. Ofei F, Hurel S, Newkirk J, Sopwith M, and Taylor R. Effects of an engineered human anti-TNF-alpha antibody (CDP571) on insulin sensitivity and glycemic control in patients with NIDDM. Diabetes. 1996;45(7):881–5.

26. Paquot N, Castillo MJ, Lefebvre PJ, and Scheen AJ. No increased insulin sensitivity after a single intravenous administration of a recombinant human tumor necrosis factor receptor: Fc fusion protein in obese insulin-resistant patients. J Clin Endocrinol Metab. 2000;85(3):1316–9.

27. Ying W, Riopel M, Bandyopadhyay G, Dong Y, Birmingham A, Seo JB, et al. Adipose Tissue Macrophage-Derived Exosomal miRNAs Can Modulate In Vivo and In Vitro Insulin Sensitivity. Cell. 2017;171(2):372–84 e12.

28. Kulkarni RN, Jhala US, Winnay JN, Krajewski S, Montminy M, and Kahn CR. PDX-1 haploinsufficiency limits the compensatory islet hyperplasia that occurs in response to insulin resistance. J Clin Invest. 2004;114(6):828–36.

29. Aamodt KI, and Powers AC. Signals in the pancreatic islet microenvironment influence beta-cell proliferation. Diabetes Obes Metab. 2017;19 Suppl 1:124–36.

30. Kulkarni RN, Mizrachi EB, Ocana AG, and Stewart AF. Human beta-cell proliferation and intracellular signaling: driving in the dark without a road map. Diabetes. 2012;61(9):2205–13.

31. Alonso LC, Yokoe T, Zhang P, Scott DK, Kim SK, O’Donnell CP, et al. Glucose infusion in mice: a new model to induce beta-cell replication. Diabetes. 2007;56(7):1792–801.

32. Porat S, Weinberg-Corem N, Tornovsky-Babaey S, Schyr-Ben-Haroush R, Hija A, Stolovich-Rain M, et al. Control of pancreatic beta cell regeneration by glucose metabolism. Cell Metab. 2011;13(4):440–9.

33. Garcia-Ocana A, Takane KK, Syed MA, Philbrick WM, Vasavada RC, and Stewart AF. Hepatocyte growth factor overexpression in the islet of transgenic mice increases beta cell proliferation, enhances islet mass, and induces mild hypoglycemia. J Biol Chem. 2000;275(2):1226–32.

34. Araujo TG, Oliveira AG, Carvalho BM, Guadagnini D, Protzek AO, Carvalheira JB, et al. Hepatocyte growth factor plays a key role in insulin resistance-associated compensatory mechanisms. Endocrinology. 2012;153(12):5760–9.

35. Assmann A, Ueki K, Winnay JN, Kadowaki T, and Kulkarni RN. Glucose effects on beta-cell growth and survival require activation of insulin receptors and insulin receptor substrate 2. Mol Cell Biol. 2009;29(11):3219–28.

36. Ying W, Lee YS, Dong Y, Seidman JS, Yang M, Isaac R, et al. Expansion of Islet-Resident Macrophages Leads to Inflammation Affecting beta Cell Proliferation and Function in Obesity. Cell Metab. 2019;29(2):457–74 e5.

37. Ying W, Fu W, Lee YS, and Olefsky JM. The role of macrophages in obesity-associated islet inflammation and beta-cell abnormalities. Nat Rev Endocrinol. 2020;16(2):81–90.

38. Filios SR, and Shalev A. beta-Cell MicroRNAs: Small but Powerful. Diabetes. 2015;64(11):3631–44.

39. Zhu M, Wei Y, Geissler C, Abschlag K, Corbalan Campos J, Hristov M, et al. Hyperlipidemia-Induced MicroRNA-155-5p Improves beta-Cell Function by Targeting Mafb. Diabetes. 2017;66(12):3072–84.

40. Ying W, Cheruku PS, Bazer FW, Safe SH, and Zhou B. Investigation of macrophage polarization using bone marrow derived macrophages. J Vis Exp. 2013(76).

41. Yuyama K, Sun H, Mitsutake S, and Igarashi Y. Sphingolipid-modulated exosome secretion promotes clearance of amyloid-beta by microglia. J Biol Chem. 2012;287(14):10977–89.

42. Bartel DP. MicroRNAs: genomics, biogenesis, mechanism, and function. Cell. 2004;116(2):281–97.

43. O’Connell RM, Taganov KD, Boldin MP, Cheng G, and Baltimore D. MicroRNA-155 is induced during the macrophage inflammatory response. Proc Natl Acad Sci U S A. 2007;104(5):1604–9.

44. Guay C, Kruit JK, Rome S, Menoud V, Mulder NL, Jurdzinski A, et al. Lymphocyte-Derived Exosomal MicroRNAs Promote Pancreatic beta Cell Death and May Contribute to Type 1 Diabetes Development. Cell Metab. 2019;29(2):348–61 e6.

45. Crewe C, Joffin N, Rutkowski JM, Kim M, Zhang F, Towler DA, et al. An Endothelial-to-Adipocyte Extracellular Vesicle Axis Governed by Metabolic State. Cell. 2018;175(3):695–708 e13.

46. Mathieu M, Martin-Jaular L, Lavieu G, and Thery C. Specificities of secretion and uptake of exosomes and other extracellular vesicles for cell-to-cell communication. Nat Cell Biol. 2019;21(1):9–17.

47. Gupta D, Jetton TL, LaRock K, Monga N, Satish B, Lausier J, et al. Temporal characterization of beta cell-adaptive and -maladaptive mechanisms during chronic high-fat feeding in C57BL/6NTac mice. J Biol Chem. 2017;292(30):12449–59.

48. Hull RL, Kodama K, Utzschneider KM, Carr DB, Prigeon RL, and Kahn SE. Dietary-fat-induced obesity in mice results in beta cell hyperplasia but not increased insulin release: evidence for specificity of impaired beta cell adaptation. Diabetologia. 2005;48(7):1350–8.

49. Chen Y, Siegel F, Kipschull S, Haas B, Frohlich H, Meister G, et al. miR-155 regulates differentiation of brown and beige adipocytes via a bistable circuit. Nat Commun. 2013;4:1769.

50. Hsin JP, Lu Y, Loeb GB, Leslie CS, and Rudensky AY. The effect of cellular context on miR-155-mediated gene regulation in four major immune cell types. Nat Immunol. 2018;19(10):1137–45.

