## Supplemental figures for "Adipose tissue macrophages orchestrate β cell adaptation in obesity through secreting miRNA-containing extracellular vesicles"

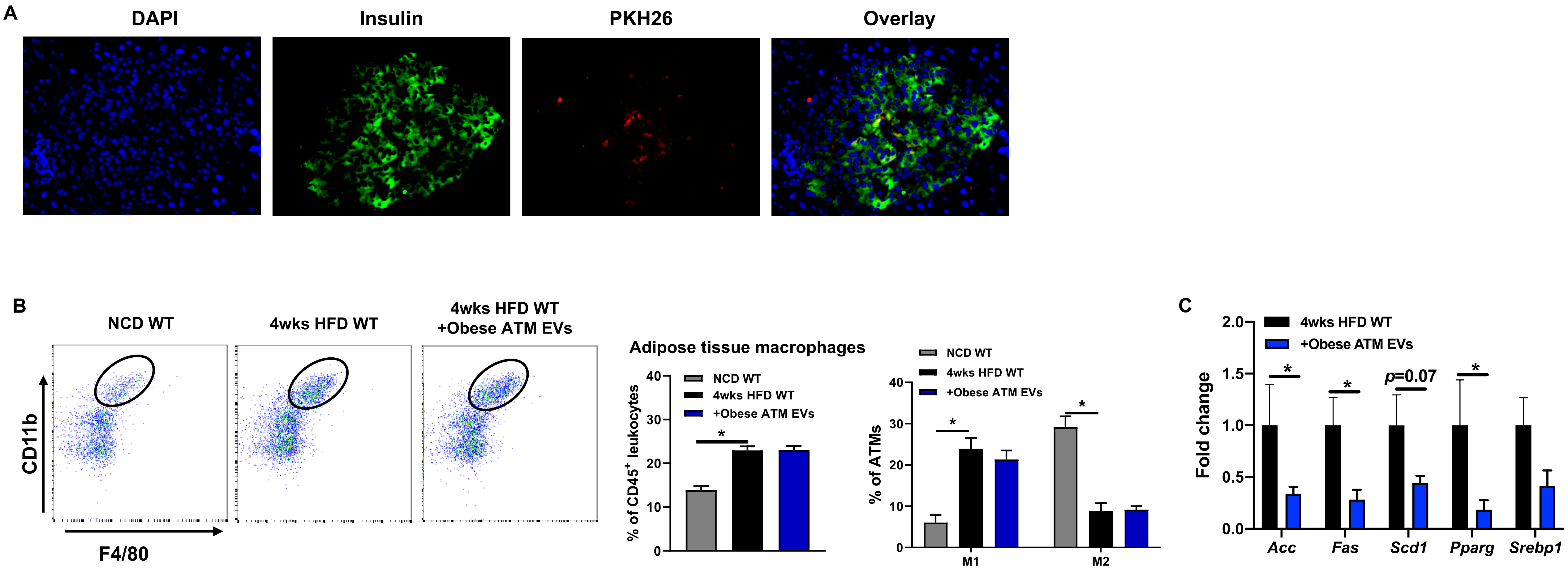


**Supplementary Figure 1. The effects of obese ATM EVs on HFD WT mice.** (A) The appearance of red fluorescence in the islets of recipients after 24 hours administration of PKH26-labeled obese ATM EVs. Representative images are shown from 3 independent experiments. (B) The population and activation of ATMs after 4 weeks treatment of obese ATM EVs. ATMs are CD45+CD11b+F4/80+ cells, and M1 and M2 ATMs were identified by CD11c+CD206- and CD11c-CD206+, respectively. (C) The expression of genes associated with lipogenesis of epididymal fat after 4 weeks treatment of obese ATM EVs. Data are presented as the mean ± SEM. B and C, n=4 per group. * $P$<0.05, Student's t test.

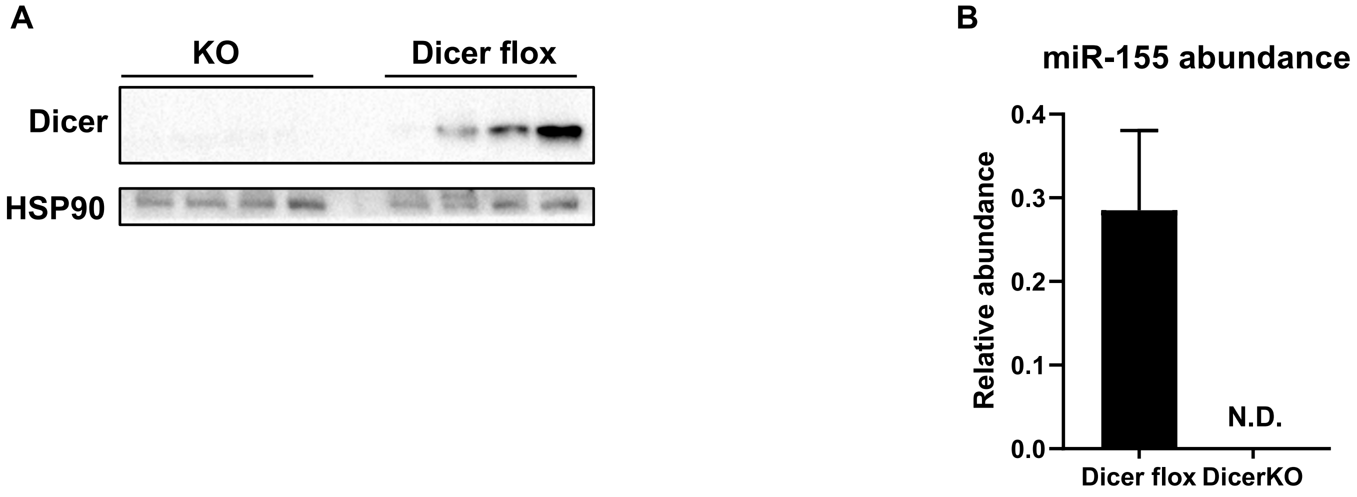


**Supplementary Figure 2. Validation of dicer knockout in obese ATMs.** (A) The abundance of dicer in ATMs isolated from 16wks HFD-fed LysMcre-Dicer flox (KO) or Dicer flox mice. (B) miR-155 abundance in ATMs after knockout of dicer. Data are presented as the mean ± SEM. B, n=3 per group. N.D., non-detectable.

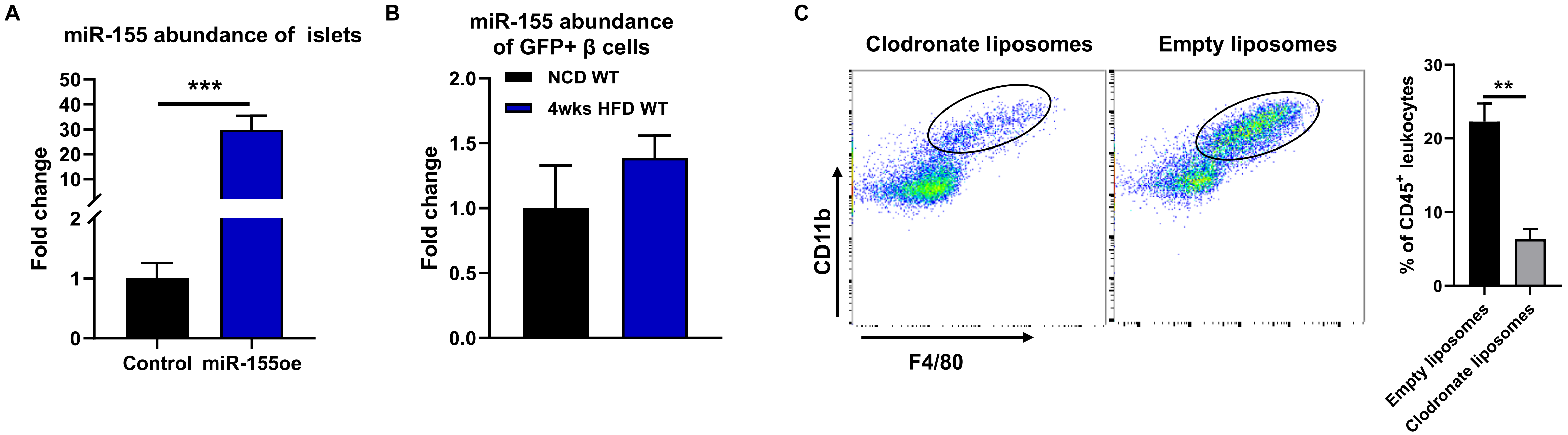


**Supplementary Figure 3. Effects of miR-155 on β cell functions.** (A) miR-155 abundance in islets after 24 hours transfection of miR-155 mimics. (B) The expression of miR-155 in GFP+ cells after 4 weeks HFD feeding. (C) The ATM population of HFD WT mice after treatment of either empty liposomes or clodronate liposomes. Data are presented as the mean ± SEM. A-C, n=4 per group. * *P*<0.05, ** *P*<0.01, ** *P*<0.001, Student's t test.

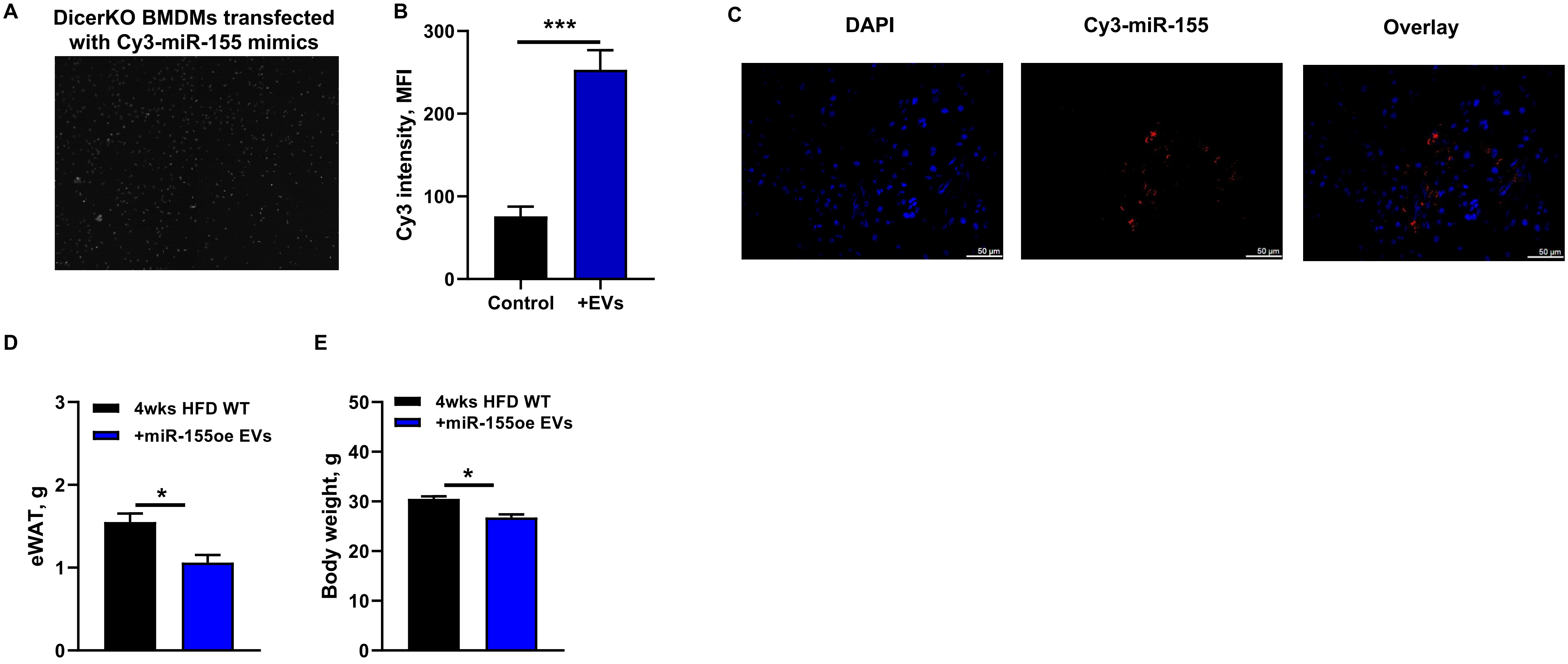


**Supplementary Figure 4. The critical role of miR-155 in macrophage-derived EVs.** (A) The presence of Cy3 fluorescent signals in DicerKO BMDMs after 24 hours transfection of Cy3-labeled miR-155 mimics. Representative images are shown from 3 independent experiments. (B) The Cy3 intensity within BMDM-derived EVs after transfection of Cy3-miR-155 mimics. (C) The appearance of Cy3 red fluorescence in the pancreas of obese recipient mice after 24 hours injection of Cy3-miR-155 containing BMDM EVs. Representative images are shown from 3 independent experiments. (D and E) The epididymal fat mass and body weight of recipients after 4 weeks treatment of miR-155oe EVs. Data are presented as the mean ± SEM. B, n=3 per group; D and E, n=5 per group. * *P*<0.05, *** *P*<0.001, Student's t test.

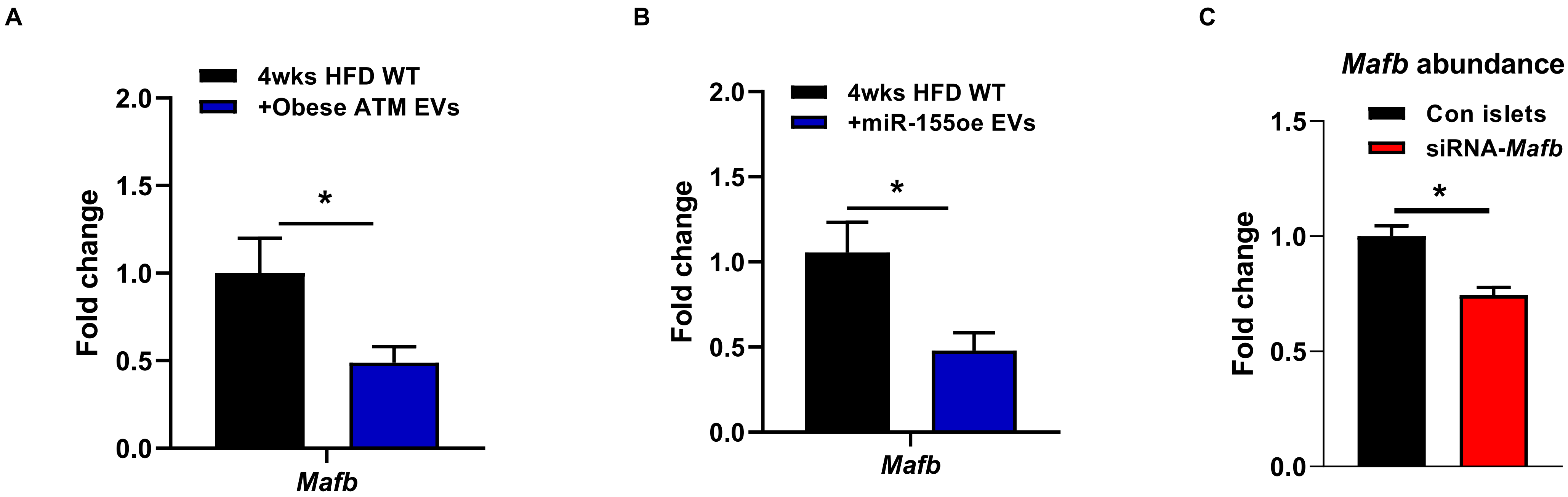


**Supplementary Figure 5. The effect of *Mafb* on β cell functions.** (A and B) The expression of *Mafb* in GFP+ cells of recipients treated with either obese ATM EVs or miR-155oe EVs for 4 weeks. (C) *Mafb* abundance in NCD islets after transfection of siRNA-*Mafb*. Data are presented as the mean ± SEM. A-C, n=4 per group. * *P*<0.05, Student's t test.
